# Sound transmission by chamber prosthesis of the middle ear

**DOI:** 10.1101/2020.10.23.352245

**Authors:** Wiktor L. Gambin

## Abstract

Tests done on specimens cut from the temporal bones show, that the stapedotomy can be more effective, if instead of the piston prosthesis, the ear chamber prosthesis is used. In that case, the vibrations of the eardrum are transferred to a plate with attachment sticked to the incus. The plate is suspended on a membrane stretched on the base of conical chamber which is filled with a fluid and placed in the middle ear cave. The sound wave caused by a vibrating plate, is focused at the chamber outlet placed in a small hole drilled in the stapes footplate. As in the case of the piston prosthesis behavior of the round window membrane differs from that observed in the normal ear. The flow through a narrow outlet of the conical chamber makes a more deflection of the central part of the round window membrane. The properties of the prosthesis elements are close to those of the removed parts of the middle ear. In spite of this, one can observe a different sound transmission inside the ear. When the sound is higher than 1000 Hz, the vibration amplitude of the plate is 5-10 dB higher than that for the stapes footplate in the healthy ear. However, when the sound is lower than 1000 Hz, this amplitude is lower than that for the stapes footplate. To explain it, a simplified model of the sound propagation in the ear given in the prior work is used. To get a better agreement with the test results, the model takes into account a damping of the sound wave by the round window membrane. Next, the model is adapted to the ear with chamber prosthesis. The factors that may have an effect on the behavior of the sound wave are examined. The first is shortening of the incus. It increases the leverage of the ossicles and the force acting on the prosthesis plate compared to that in the normal ear. Next factor is a reduction of the mass of the vibrating plate what makes a growth of its resonance frequency. This slightly reduces the amplitude of the plate for the low sounds and increases it for the medium and the higher sounds. At end, the lack of the influence of the flow through the conical chamber on the sound wave energy is shown. The assumed model gives the rules for amplitudes of the chamber plate as functions of the sound frequency. Their values for the sound frequency from 400 Hz to 8000 Hz and its graphs are shown and compared with those for the stapes footplate in the normal ear. One can see that if the sound frequency is higher than 1000 Hz, then the chamber prosthesis makes higher amplitudes of the sound wave than the normal ear. To explain their drop for frequencies lower than 1000 Hz, needs more tests in this range.

## 1. Introduction

A commonly used implant of the middle ear is the piston prosthesis. It improves the hearing of patients in the range of medium sound frequencies, and thus it enables a speech perception. However, the high and low sounds are not recognized enough. Why this is so and how to improve the efficiency of the piston prosthesis is discussed in [1].

Another way to increase the frequency range of the heard sounds is to use the chamber prosthesis [2–4]. The main part of this prosthesis is a truncated cone-shaped chamber filled with a fluid and ending in a thin capillary tube with a length of 1 mm (Fig. 1). In place of the cut off stapes legs, the chamber with a height of 3 mm is placed, and its outlet tube is placed in a hole with a diameter 0.6 mm drilled in the stapes foot. The inlet to the chamber is closed by a rigid plate stuck to the incus long process and suspended on a thin membrane. The sound wave caused by the vibrations of the plate focuses on the outlet of the chamber, then expands inside the cochlea and as a plane wave reaches the round window (RW). The chamber and rigid plate are made from ABS plastic with 3D rapid prototyping technique. The flexible membrane is made from the liquid UV-light cured adhesive NOA 68.

**Fig. 1.**
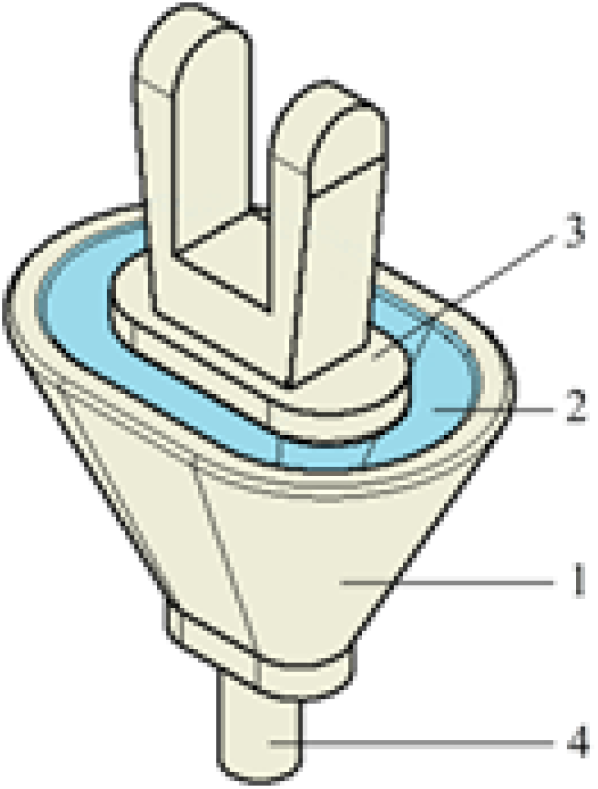
Chamber prosthesis: 1 – conical chamber filled with fluid, 2 – flexible membrane, 3 – rigid plate with an attachment to the incus long process, 4 – thin capillary tube; (from [4]).

The properties and dimensions of the prosthesis and the removed parts of the ear, which will be used, are listed in the Table 1. The data is from the work [4] and a project done by Dr. M. Sołyga from Warsaw University of Technology.

**Table 1.**

| Element & property |  | Chamber prosthesis | Middle ear |
| --- | --- | --- | --- |
| Plate & attachment /<br>Stapes | Surface area | $F_{ch} = 2.8 \text{ [mm}^2\text{]}$ | $F_{SF} = 2.5 \text{ [mm}^2\text{]}$ |
| | Mass | $m_{ch}^{*)} = 1.50 \text{ [mg]}$ | $m_{SF} = 3.04 \text{ [mg]}$ |
| | Natural frequency | $f_{ch} = \frac{1}{2\pi} \sqrt{\frac{R_{ch}}{m_{ch}}} = 1500 \text{ [s}^{-1}\text{]}$ | $f_{SF} = \frac{1}{2\pi} \sqrt{\frac{R_{SF}}{m_{SF}}} = 1000 \text{ [s}^{-1}\text{]}$ |
| Membrane / angular<br>ligament | Stiffness | $R_{ch}^{**}) = 0.134 \text{ [N/mm]}$ | $R_{SF} = 0.120 \text{ [N/mm}^2\text{]}$ |
| Chamber | Basis area | $F_{ch}^0 = 6.7 \text{ [mm}^2\text{]}$ | ————— |
| | Outlet area | $F_{ch}^1 = 0.07 \text{ [mm}^2\text{]}$ | |

One can see that all parts of the prosthesis imitate damaged or removed parts of the ear. In spite of this, there appear three new effects are seen in the normal ear. First, when the sound is higher than 1000Hz, the vibration amplitude of the plate is 5-10 dB higher than that for the stapes footplate in the healthy ear. In the work [4], it was introduced as a logarithmic measure of the difference of the vibration amplitude of the plate and the stapes footplate. The measure is called the Input Ratio (dB), and its definition for the piston prosthesis and the chamber prosthesis can be found at Fig. 2. There a comparison of the Input Ratio values for the both prostheses is shown too.

**Fig. 2.**
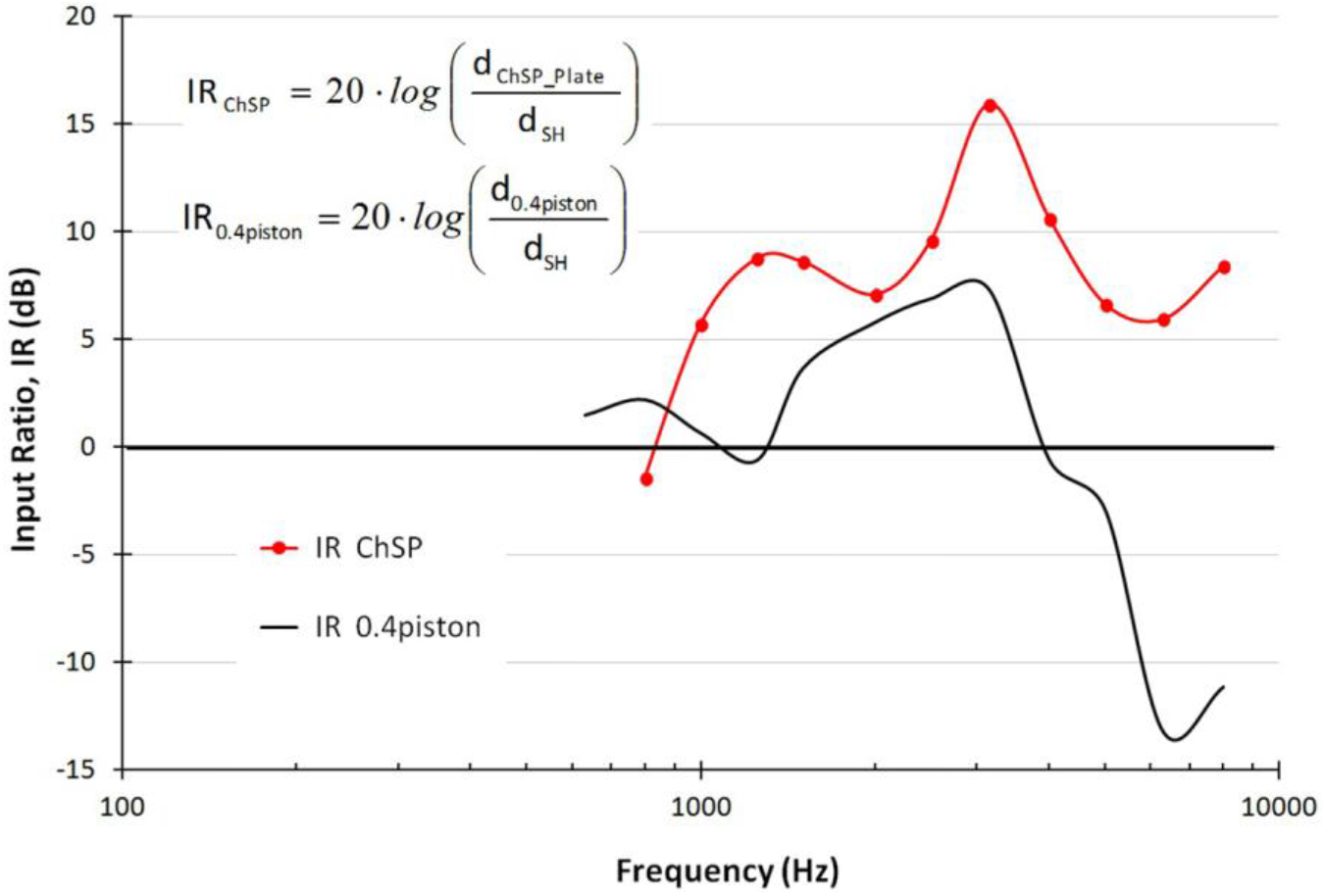
Input Ratio-frequency characteristics: the chamber prosthesis – red line, the 0.4-mm piston prosthesis - black line; (from [4]).

The second effect appears when the sound is lower than 1000 Hz. Then the vibration amplitude of the plate is lower than that for the stapes footplate as it is shown at Fig. 2. This effect does not appear in the case of the piston prosthesis. But the rounding of the wave front caused by the thin end of the piston makes a reduction in the acoustic pressure acting on the basilar membrane. As a result, the both prostheses give a similar perception of the low sounds.

At last, one can observe that, unlike in the normal ear, a small central part of the circular window membrane bulges much more than the rest of the membrane (Fig.3). The same effect one can see for the piston prosthesis, but for the low sounds only. For the higher sounds there is a noise effect caused by a secondary wave described in [1].

**Fig. 3.**
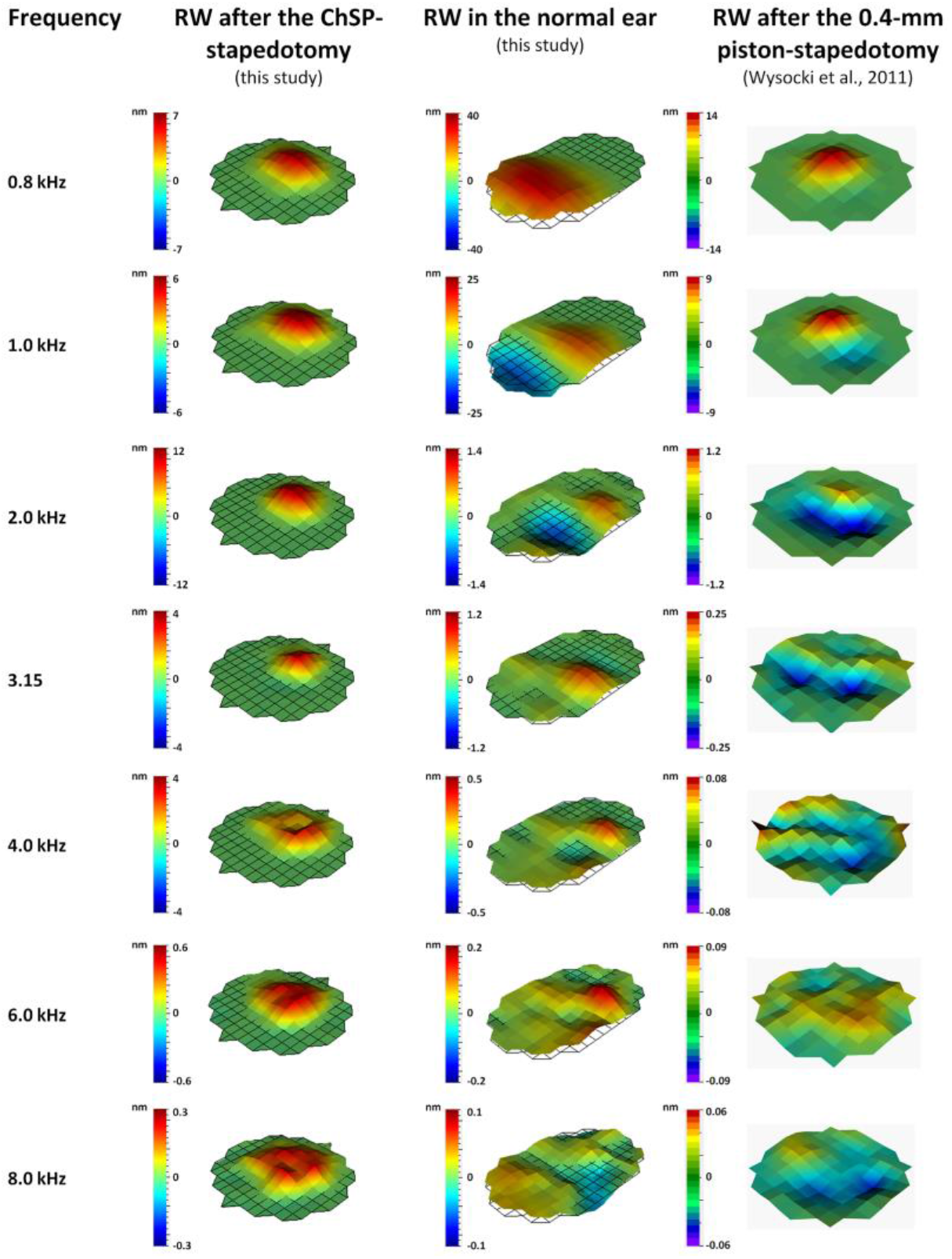
Three-dimensional visualization of displacements of the round window membrane after stapedotomy with chamber prosthesis (left), 0.4-mm piston prosthesis (right) and normal ear (middle); (from [4]).

To find a cause of these effects, a way of the sound wave propagation in the ear with the chamber prosthesis is compared with that in the normal ear. For this purpose, a simplified model of sound propagation in the human ear proposed and tested in the work [5] is used. The model gives rules for the amplitudes of the stapes-footplate and of the round window in the healthy ear, if the sound source frequency and its intensity are known. Also rules for amplitudes of points of the basilar membrane are given. It enables to state a level of the cochlear amplification of the sound in the ear. The model does not take into account a dissipation of the sound energy in the cochlea. It gives that the amplitude of the stapes footplate for the test frequency 3150 Hz is too large. Here, the model is changed by taking into account a loss of the sound energy due to viscous properties of the round window membrane. Next, these rules taken out from the upgraded model are suited to the ear with the chamber prosthesis.

## 2. Upgrade of the simplified model

### 2.1 Comparison of results for the normal ear

In the work [5], for the sound intensity level ***β*** = **90 *dB***, the amplitudes of the stapes footplate *A*_*SF*_ are given and compared with those measured in [3] shown here as 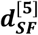. Let us explain what makes differences between them. The presented model bases on the assumed dimensions of parts of the ear. The force ***N***_***SF***_ acting on the stapes footplate depends only on the cross-sectional area of the inlet to the ear canal. If its area is ***F***_**0**_, then

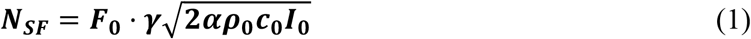

where ***γ*** is the malleus-incus lever ratio, ***α*** – the absorption coefficient of the sound wave, ***ρ***_**0**_ – the air density, ***c*_0_** – the sound velocity in air (at 20° C) and ***I***_**0**_ is the given sound intensity. The static displacement of the stapes footplate due to this force is

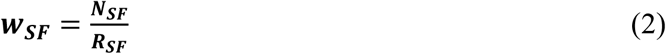

where 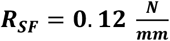 is stiffness of the annular ligament. Because the resonant frequency of the stapes is **1 *kHz***, then ***w***_***SF***_ should be equal to the measured amplitude 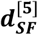 for ***f***_**(*i*)**_ = **1*kHz***. It was taken ***F***_**0**_ = **3 *mm***, in the work [5], and it has been obtained ***w***_***SF***_ = **1.17 · 10**^**−5**^ *mm*. A value of the measured amplitude given in [3] is 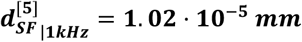. Notice that the measured and calculated amplitudes would be the same if the inlet area would be 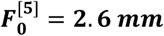. Here, the results for the normal ear and the ear with the chamber prosthesis obtained from the simplified model are compared with the test results given in [4]. From [4], the measured amplitude is 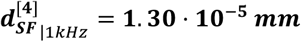 for the normal ear. It suits to the inlet area 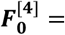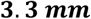. Notice that the tests on temporal bone samples do not let take into account a real size of the area of the inlet to the ear canal (Fig. 4a,b). Also, one can find that the differences between the measured and calculated amplitudes are of the same order. That is why ***we take an average inlet area F***_**0**_ = **3 *mm***. Then the static displacement of the stapes footplate and the corresponding force are taken as

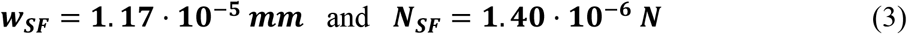

**Fig. 4.**
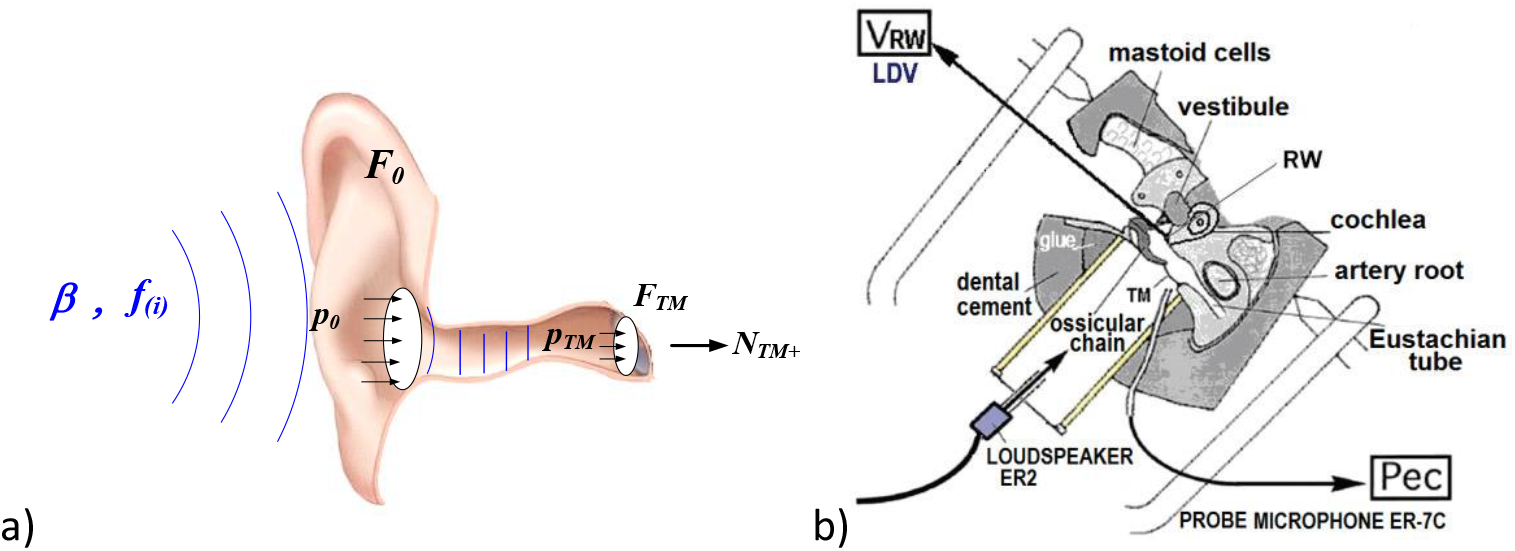
Sound wave propagation in the outer ear: a) in the model (from [5]), b) in the tests (from [6]).

According to the assumed model, the dynamic amplitude of the stapes footplate ***A*_*SF*_** is determined by ***w***_***SF***_ by the rule

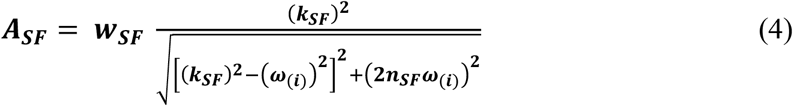

where ***ω***_(***i***)_ = **2π · *f***_(***i***)_ is the given angular frequency of the sound, 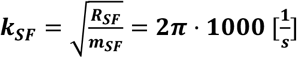 is the angular resonant frequency of the stapes and **2*n***_***SF***_ = ***k***_***SF***_ is the damping factor.

As measure of the difference between the measured 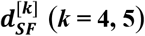 and the calculated ***A***_***SF***_ values, one can take

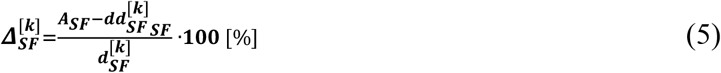

The values of ***A***_***SF***_, 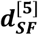 and 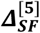 given in [5], as well as, and the values of ***A***_***SF***_, 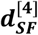 and 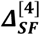 used in this work are shown in Table 2. At the last column of Table 2, relative differences between two tests 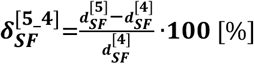 are given.

**Table 2.**

| Test<br>$t$<br>$(i)$ | Frequency<br>$f_{(i)}$ [Hz] | Calculated<br>amplitude<br>$A_{SF}$ [mm] | Measured<br>amplitude<br>(from [5])<br>$d_{SF}^{[5]}$ [mm] | Relative<br>difference<br>$\Delta_{SF}^{[5]}$ [%] | Measured<br>amplitude<br>(from [4])<br>$d_{SF}^{[4]}$ [mm] | Relative<br>difference<br>$\Delta_{SF}^{[4]}$ [%] | Tests<br>difference<br>$\delta_{SF}^{[5,4]}$ [%] |
| --- | --- | --- | --- | --- | --- | --- | --- |
| 1 | 400 | $1.2576 \cdot 10^{-05}$ | $1.64 \cdot 10^{-05}$ | -23 | - | - | - |
| 2 | 500 | $1.2980 \cdot 10^{-05}$ | $1.53 \cdot 10^{-05}$ | -15 | - | - | - |
| 3 | 630 | $1.3415 \cdot 10^{-05}$ | $1.90 \cdot 10^{-05}$ | -29 | - | - | - |
| 4 | 800 | $1.3337 \cdot 10^{-05}$ | $1.97 \cdot 10^{-05}$ | -32 | $2.81 \cdot 10^{-5}$ | -52 | -30 |
| 5 | 1000 | $1.1700 \cdot 10^{-05}$ | $1.02 \cdot 10^{-05}$ | 15 | $1.30 \cdot 10^{-5}$ | -10 | -22 |
| 6 | 1250 | $8.5351 \cdot 10^{-06}$ | $8.92 \cdot 10^{-06}$ | -4 | $7.52 \cdot 10^{-6}$ | 13 | 19 |
| 7 | 1500 | $5.9921 \cdot 10^{-06}$ | - | - | $6.10 \cdot 10^{-6}$ | -2 | - |
| 8 | 1600 | $5.2354 \cdot 10^{-06}$ | $7.12 \cdot 10^{-06}$ | -26 | - | - | - |
| 9 | 2000 | $3.2448 \cdot 10^{-06}$ | $4.93 \cdot 10^{-06}$ | -34 | $4.07 \cdot 10^{-6}$ | -20 | 21 |
| 10 | 2500 | $2.0120 \cdot 10^{-06}$ | $2.84 \cdot 10^{-06}$ | -29 | $2.02 \cdot 10^{-6}$ | 0 | 41 |
| 11 | 3150 | $1.2364 \cdot 10^{-06}$ | $6.40 \cdot 10^{-07}$ | 93 | $6.04 \cdot 10^{-7}$ | 105 | 6 |
| 12 | 4000 | $7.5362 \cdot 10^{-07}$ | $7.73 \cdot 10^{-07}$ | -3 | $6.42 \cdot 10^{-7}$ | 17 | 21 |
| 13 | 5000 | $4.7722 \cdot 10^{-07}$ | $8.36 \cdot 10^{-07}$ | -43 | $7.03 \cdot 10^{-7}$ | -32 | 19 |
| 14 | 6000 | $3.2947 \cdot 10^{-07}$ | - | - | $5.51 \cdot 10^{-7}$ | -40 | - |
| 15 | 6300 | $2.9845 \cdot 10^{-07}$ | $6.14 \cdot 10^{-07}$ | -51 | - | - | - |
| 16 | 8000 | $1.8422 \cdot 10^{-07}$ | $1.69 \cdot 10^{-07}$ | 9 | $1.82 \cdot 10^{-7}$ | 1 | -7 |
| 17 | 10000 | $1.1757 \cdot 10^{-07}$ | $7.57 \cdot 10^{-08}$ | 55 | - | - | - |

Let us pay attention to the differences between the test values 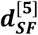 and ***d*_*SF*_**. Note that the highest absolute value of these differences is **41%**. On the other hand, apart the frequency of **3150 Hz**, the difference between 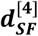 and ***A***_***SF***_ which yield from the model does not surpass **52%**. For the frequency of **3150 Hz**, a strong damping is observed in both tests, which is not included in the model proposed in [5]. For this frequency, the difference between 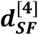 and ***A***_***SF***_ go up to **105%**. To reduce this difference, a new assumption should be made to the model.

### 2.2 A modification of the model

Recall that for the frequency of **3150 *Hz***, both tests show a two times lower amplitudes than it yields from the model. This is the frequency at which ***equal loudness contours***, known as Fletcher-Munson curves reach a minimum [7] (see Fig. 6). This means that the human ear can hear sounds at this frequency best, in spite of the fact that tests on samples of the temporal bones then show strong damping.

**Fig. 5.**
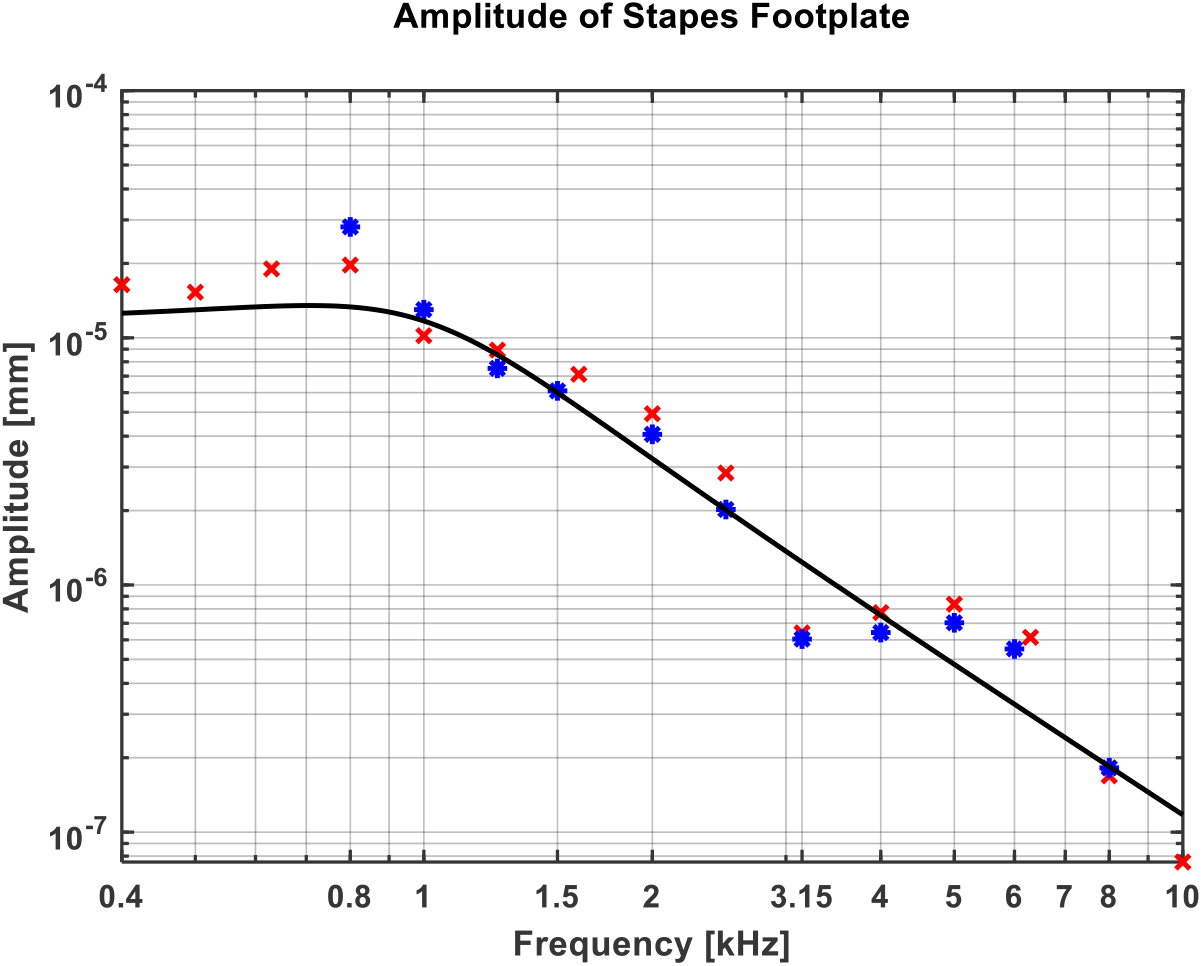
Amplitude of stapes footplate: measured (x – from [5] and * – from [4]) and calculated (solid line).

**Fig. 6.**
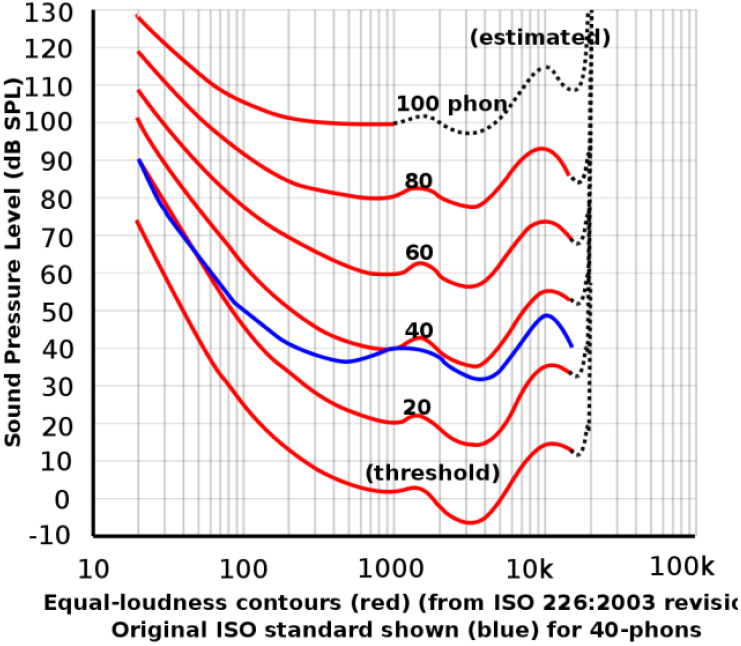
Equal-loudness contours (from ISO 226:2003).

To explain this contradiction, one should take into account the behavior of the sound wave in the ear canal before it reaches the eardrum. Similar to wind instruments, reflections from the walls of the ear canal cause the standing wave. This wave gives a resonance at the specific sound frequency ***f***_**0**_. If ***c* = 340 *m/s*** is sound velocity in air and ***L* = 0. 027 *m***; is the length of the auditory canal, then the resonance frequency is

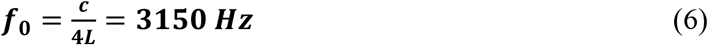

The resonance of the sound wave in the ear canal causes that the ear is the most sensitive to the frequency of **3150 *Hz***. This resonance does not appear during the tests on the temporal bone samples. Instead of that, some amplification of low (***f***_(***i***)_ ≤ **800 *Hz***) and high (***f***_(***i***)_ ≥ **5000 *Hz***) sounds appears (see Table2). Perhaps the cause is a long sound input tube of the speaker adapter (see Fig. 4b). Note that 12 cm long tube makes a standing wave with the amplitude 700 Hz, and the length of 12 mm makes its tenth harmonic with the amplitude 7000 Hz.

Contrary to the above hearing tests, one can observe strong sound damping during tests on samples of the temporal bone for the frequency **3150 *Hz***. The main reason of this fact is the damping of the sound wave in the cochlea due to the ***inelastic behavior of the round window membrane***. Up to now, our model assumes that there is no the sound wave energy dissipation. In fact, such dissipation takes place (see [8]). Let ***P***_***SF***_ is the input power of the stapes footplate. Denote by ***D***_***RW***_, a part of ***P***_***SF***_ absorbed by vibrations of the round window membrane. Then the real power of the stapes footplate is

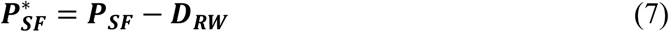

Following the work [8], let us introduce the *loss factor* 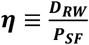. Now, Eq. (7) takes the form

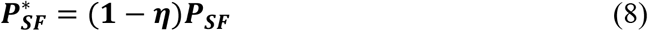

According to [5], for a given ***A***_***SF***_, the input power of the stapes footplate is

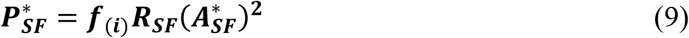

where, the reduced amplitude of the stapes footplate 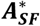 is

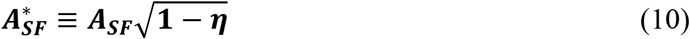

The measured values of the loss factor ***η*** one can find in the paper [8] (Fig. 7a). One can approximate the above data by the function (see Fig.7b)

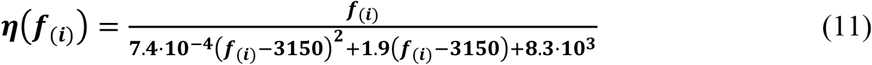

where ***f***_(***i***)_ are given in [***Hz***] or by function

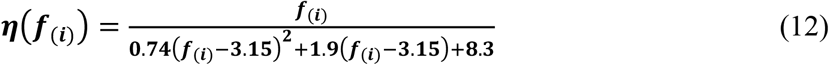

if ***f***_(***i***)_ are taken in [***kHz***]. And so, the amplitude of the stapes footplate 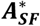 in the normal ear, given by Eq. (4), now takes the form

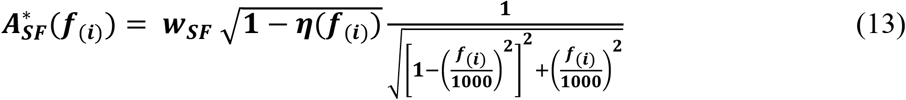

**Fig. 7.**
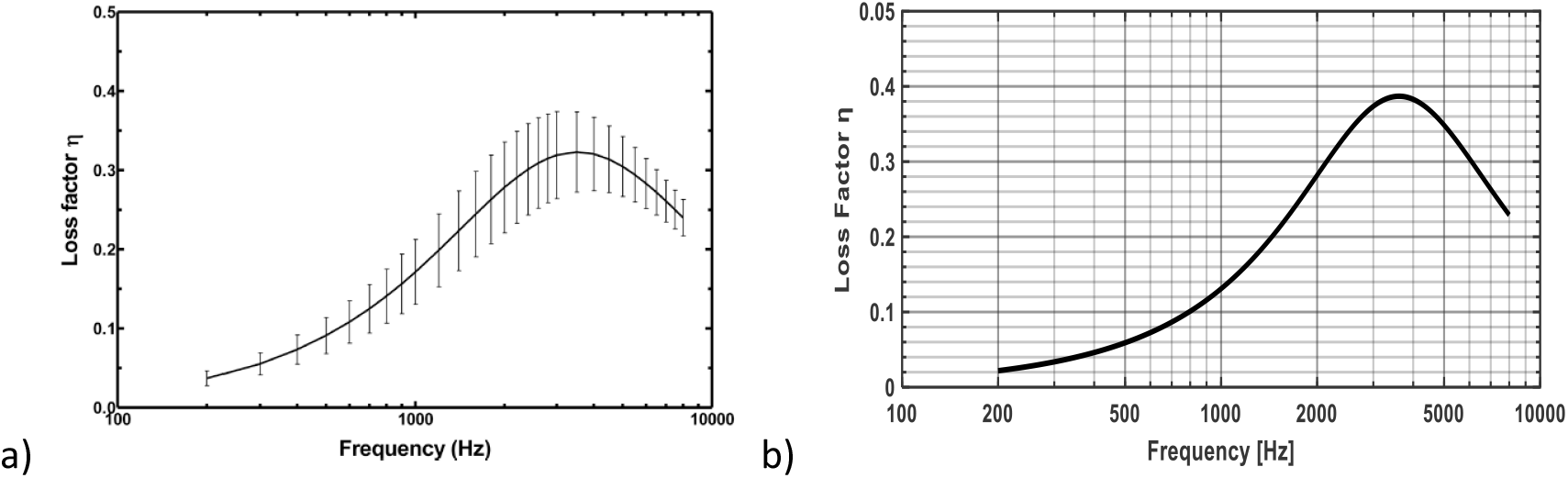
Loss factor: a) η -according to paper [8]), b) ***η***(***f***_(***i***)_) - according to Eq. (12).

**Fig. 8.**
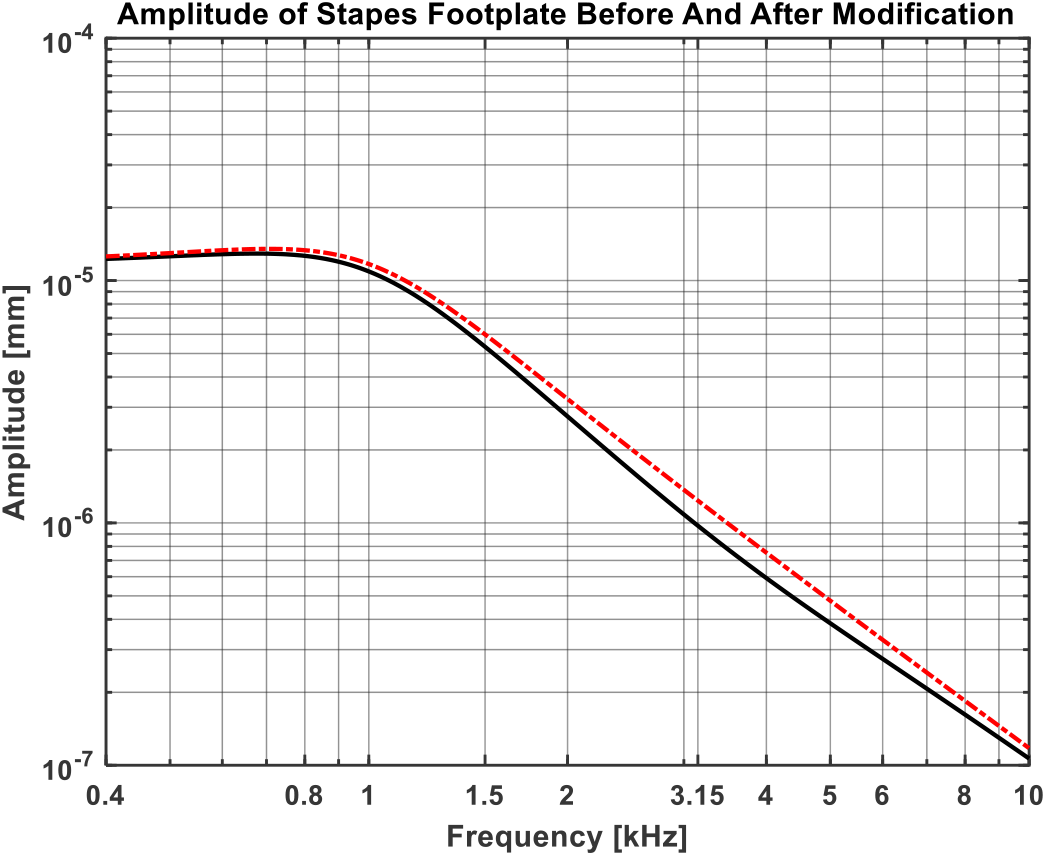
Calculated amplitudes of the stapes footplate: according to model given in [5] (dash-dot red line) and after modification (solid black line).

The results based on the modified and original model are shown in Table 3. Note that now the highest difference is **61%** for the frequency **3150 *Hz***. It means that the damping at this frequency is higher than shown in [8], or may also be influenced by another factor. On the other hand, the highest difference between the test results ***d***^**[5]**^ and ***d***^**[4]**^ is **41%** (see Table 1).

**Table 3.**

| Test<br>(i) | Frequency<br>$f_{(i)}$ [Hz] | Calculated<br>amplitude<br>$A_{SF}$ [mm] | Measured<br>amplitude<br>$d_{SF}^{[4]}$ [mm] | Relative<br>difference<br>$\Delta_{SF}^{[4]}$ [%] | Loss<br>Factor<br>$\eta$ | Modified<br>amplitude<br>$A_{SF}^*$ [mm] | Relative<br>difference<br>$\Delta_{SF}^*$ [%] |
| --- | --- | --- | --- | --- | --- | --- | --- |
| 1 | 800 | $1.3337 \cdot 10^{-5}$ | $2.81 \cdot 10^{-5}$ | -52 | 0.10 | $1.27 \cdot 10^{-5}$ | -55 |
| 2 | 1000 | $1.1700 \cdot 10^{-5}$ | $1.30 \cdot 10^{-5}$ | -10 | 0.13 | $1.09 \cdot 10^{-5}$ | -16 |
| 3 | 1250 | $8.5351 \cdot 10^{-6}$ | $7.52 \cdot 10^{-6}$ | 13 | 0.17 | $7.78 \cdot 10^{-6}$ | 3 |
| 4 | 1500 | $5.9921 \cdot 10^{-6}$ | $6.10 \cdot 10^{-6}$ | -2 | 0.21 | $5.33 \cdot 10^{-6}$ | -13 |
| 5 | 2000 | $3.2448 \cdot 10^{-6}$ | $4.07 \cdot 10^{-6}$ | -20 | 0.28 | $2.75 \cdot 10^{-6}$ | -32 |
| 4 | 2500 | $2.0120 \cdot 10^{-6}$ | $2.02 \cdot 10^{-6}$ | 0 | 0.34 | $1.63 \cdot 10^{-6}$ | -19 |
| 7 | 3150 | $1.2364 \cdot 10^{-6}$ | $6.04 \cdot 10^{-7}$ | **105** | 0.38 | $9.74 \cdot 10^{-7}$ | **61** |
| 8 | 4000 | $7.5362 \cdot 10^{-7}$ | $6.42 \cdot 10^{-7}$ | 17 | 0.38 | $5.93 \cdot 10^{-7}$ | -8 |
| 9 | 5000 | $4.7722 \cdot 10^{-7}$ | $7.03 \cdot 10^{-7}$ | -32 | 0.35 | $3.85 \cdot 10^{-7}$ | -45 |
| 10 | 6000 | $3.2947 \cdot 10^{-7}$ | $5.51 \cdot 10^{-7}$ | -40 | 0.30 | $2.76 \cdot 10^{-7}$ | -50 |
| 11 | 8000 | $1.8422 \cdot 10^{-7}$ | $1.82 \cdot 10^{-7}$ | 1 | 0.23 | $1.62 \cdot 10^{-7}$ | -11 |

## 3. The ear with the chamber prosthesis

### 3.1 Factors changing the character of the sound wave

#### 3.1.1 Change of the arm length of the malleus-incus lever

For the sound intensity ***β*** from the range from **10***dB* to **120***dB*, the values of the forces ***N***_***TM***+_ of the impact of the eardrum on the ossicles are given in [5]. Under the sound coming to the implanted chamber prosthesis, the same forces act on the prosthesis plate. In the normal ear, due the effect of the malleus-incus lever the force ***N***_***SF***_ which acts on the stapes footplate is ***N***_***SF***+_ = 1.3*N*_*TM*+_ (see Table 1 in [5], where *N*_*TM*+_ = *N*_*int*_).

Due to the cut-off of the stapes, the distance between a connection of the plate of the prosthesis with the incus and the incus head is shortened by **33%** (Fig. 9). Assume ***a*** and ***b*** as the lengths of the malleus and incus, respectively. In the normal ear, the lever ratio is ***a*: *b*** = **1.3**. If 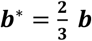 is the length of the shortened incus arm, then this ratio is ***a*: *b*\*** = **2.0**. (see Fig. 10). When ***β* = 90 *dB***, then from Eq. (3b) yields ***N***_***SF***_ = **1.40 · 10^−6^ [*N*]**. So, if ***N***_***TM***+_ = ***N***_***SF***_/**1.3** = **1.08 · 10**^**−6**^ [***N***], so

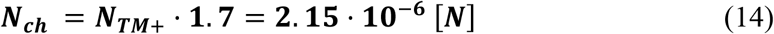

**Fig. 9.**
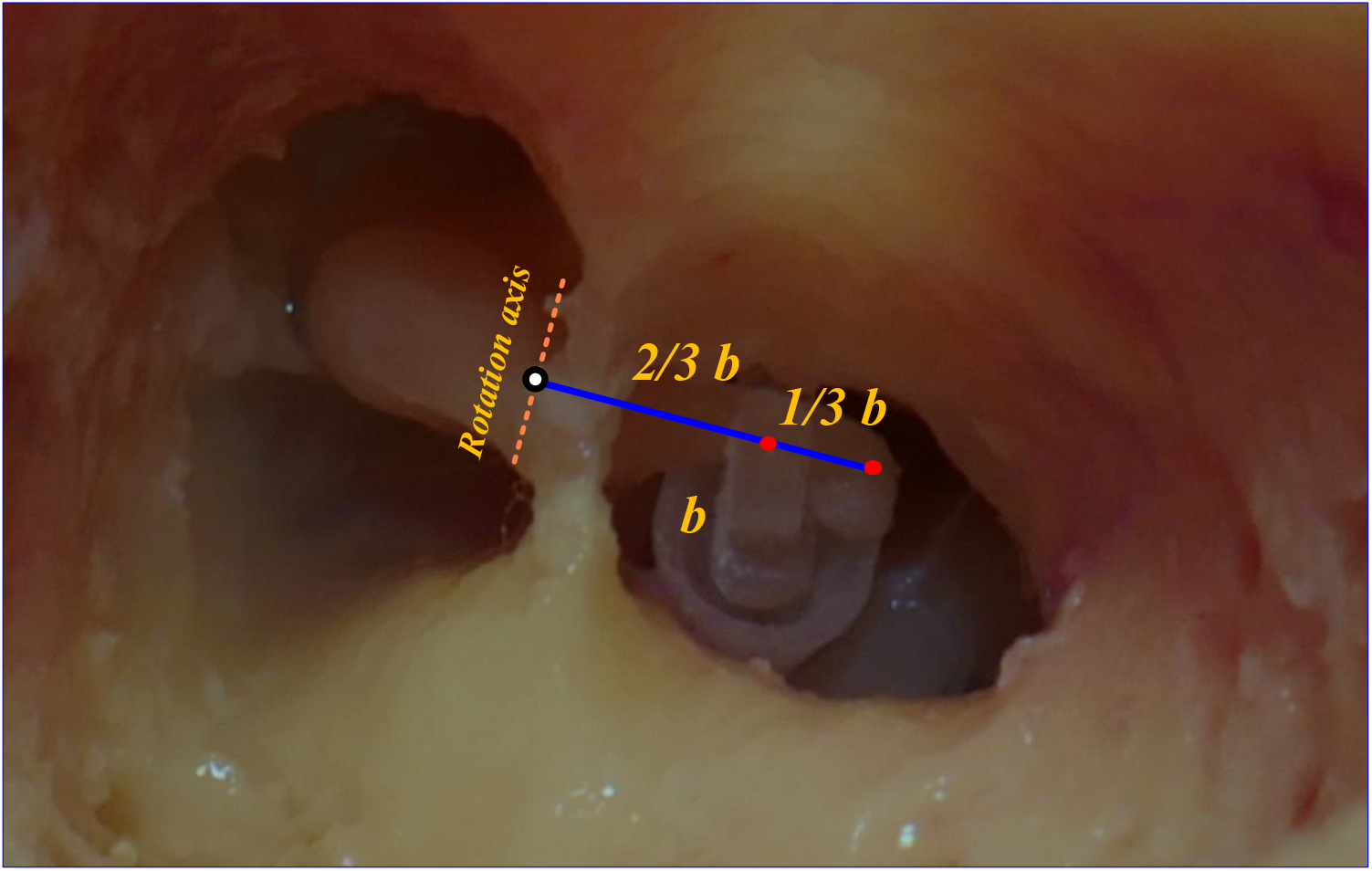
View of 20% shortening of the arm of the malleus-incus lever (from [4]).

**Fig. 10.**
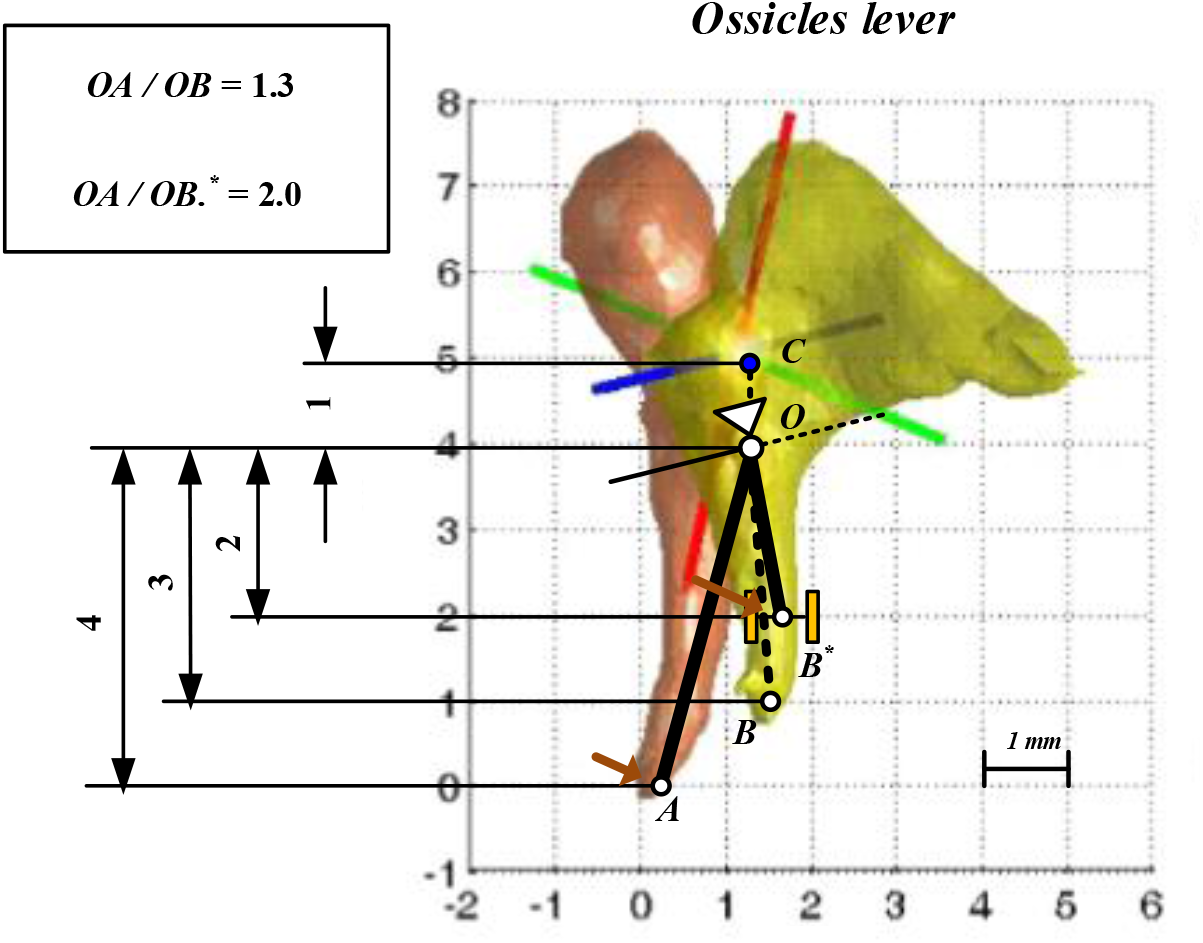
Ossicles lever ratio in the normal ear and the ear with the chamber prosthesis: ***A*** – umbo, ***B*** - incudostapedial joint, ***B*\*** - prosthesis joint, ***O*** – rotational axis, ***C*** – center of mass (based on data from [9]).

So, the force ***N***_***ch***_ acting on the plate of the prosthesis is **33%** higher than the force ***N***_***SF***_ acting on the stapes footplate in the normal ear. If ***R*_*ch*_** is the stiffness of the plate suspension (see Table 1) then the static displacement of the plate is

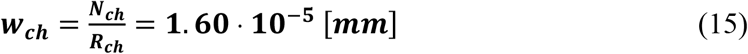

One can see that the static displacement of the plate of the prosthesis ***w*_*ch*_** is **37%** larger than the static displacement of the stapes footplate ***w*_*SF*_ = 1. 17 · 10^−5^ [*mm*]** in the normal ear.

#### 3.1.2 Mass of the plate with attachment ant its resonance frequency

In the normal ear, the mass of the stapes is assumed to determine the resonance frequency of the stapes footplate. The mass of the malleus and the incus is not taken into account.

This is due to the fact that the malleus is suspended from the walls of the middle ear niche by means of three non-collinear ligaments (**1**,**2**,**3** at Fig 11). The incus is jointed with the malleus (by ligament ***a*** at Fig. 11) and suspended from the walls of the niche through two ligaments (**4**,**5** at Fig. 11). Thus, their inertia only affects the niche walls. On the other hand, the stapes with the footplate is suspended freely from the incus. It is joined with the long process by the incudostapedial ligament (***b*** at Fig. 11) and supported by the annular ligament *c* at Fig. 11).

**Fig. 11.**
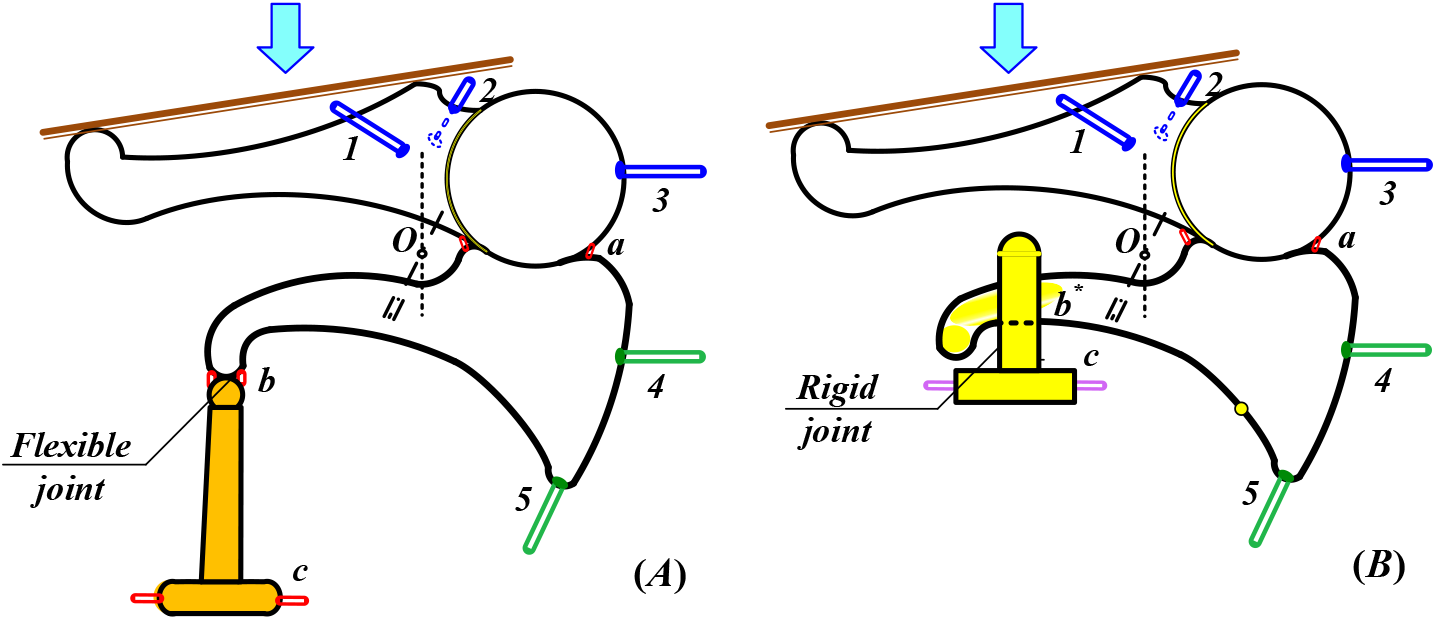
Suspension of the stapes footplate (*A*) and the prosthesis plate (*B*).

The prosthesis plate with the attachment is joined with the incus in the other way than the stapes footplate. Now, the incus arm is glued to the element making a rigid joint. Part of the mass of the plate attachment is transferred by the incus arm to the ligaments holding the incus. It is taken that two thirds of the attachment mass does not load the prosthesis membrane. Concluding, it is assumed that the mass of **1.5** *mg* acts on this membrane. One can call it the ***active mass***. On the other hand, the average value of the mass of the stapes is taken as **3.04** *mg*. The mass which load the prosthesis membrane decides on the resonant frequency of the plate, and thus o the plate amplitude. To show it, a rule for the amplitude of the prosthesis plate 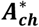 is needed. Using Eqs (4,10), one can write

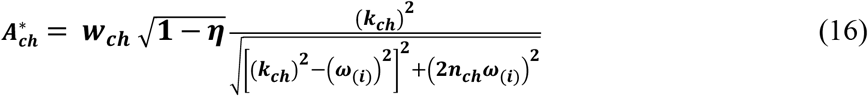

where ***η*** = ***η***(***f***_(***i***)_) is the loss factor (see Eq.(12), **ω**_(***i***)_ = **2π · *f***_(***i***)_ is the given angular frequency of the sound, 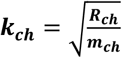 is the angular resonant frequency of the plate and **2*n***_***ch***_ is the damping factor (see [5]). The last one is given by the rule 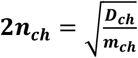. Here ***D***_***ch***_ is the parameter of the damping of the plate suspension. For the values ***R***_***ch***_ and ***m***_***ch***_ given in Table 1, we have

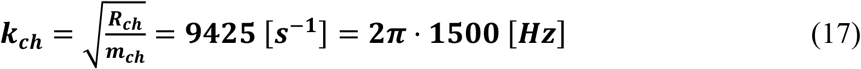

Then 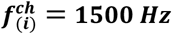 is the natural frequency of the prosthesis plate. This result is due to the ***active mass*** of the plate with the attachment, which is two times smaller than the averaged mass of the stapes.

The amplitudes 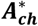 which yield from Eq. (16) are compared with the measured in [4] for frequencies **800 *Hz*** ≤ ***f***_(***i***)_ ≤ **8000 *Hz*** and shown in Table 4. Note that the obtained value of the static displacement of the plate ***w***_***ch***_ = **1. 60 · 10**^**−5**^ ***mm*** is close to the measured in [4] dynamic amplitude of the plate ***d***_***ch***_ = **1. 64 · 10**^**−5**^ ***mm*** for ***f***_(***i***)_ = **1500 *Hz***.

**Table 4.**

| *Test*<br>(i) | *Frequency*<br>$f_{(i)} [Hz]$ | *Loss*<br>*Factor*<br>$\eta$ | *Calculated*<br>*amplitude*<br>$A_{ch}^* [mm]$ | *Measured*<br>*amplitude*<br>$d_{ch} [mm]$ | *Relative*<br>*difference*<br>$\Delta_{ch}^* [\%]$ |
| --- | --- | --- | --- | --- | --- |
| 1 | 800 | 0.10 | $1.88 \cdot 10^{-5}$ | $2.34 \cdot 10^{-5}$ | -20 |
| 2 | 1000 | 0.13 | $1.90 \cdot 10^{-5}$ | $2.57 \cdot 10^{-5}$ | -26 |
| 3 | 1250 | 0.17 | $1.71 \cdot 10^{-5}$ | $2.05 \cdot 10^{-5}$ | -17 |
| 4 | 1500 | 0.21 | $1.42 \cdot 10^{-5}$ | $1.64 \cdot 10^{-5}$ | -13 |
| 5 | 2000 | 0.28 | $8.80 \cdot 10^{-6}$ | $9.06 \cdot 10^{-6}$ | -3 |
| 4 | 2500 | 0.34 | $5.33 \cdot 10^{-6}$ | $6.11 \cdot 10^{-6}$ | -9 |
| 7 | 3150 | 0.38 | $3.15 \cdot 10^{-6}$ | $3.75 \cdot 10^{-6}$ | -16 |
| 8 | 4000 | 0.38 | $1.78 \cdot 10^{-6}$ | $2.18 \cdot 10^{-6}$ | -18 |
| 9 | 5000 | 0.35 | $1.16 \cdot 10^{-6}$ | $1.50 \cdot 10^{-6}$ | -23 |
| 10 | 6000 | 0.30 | $8.37 \cdot 10^{-7}$ | $1.09 \cdot 10^{-6}$ | -23 |
| 11 | 8000 | 0.23 | $4.94 \cdot 10^{-7}$ | $4.74 \cdot 10^{-7}$ | 4 |

To get **2*n*_*ch*_**, one can take the same assumption as in the case of the normal ear that the resonant amplitude 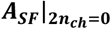 is equal to the static displacement ***w*_*SF*_** [5]. Doing it, we get 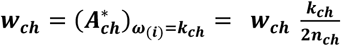 and then

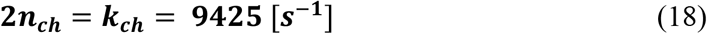

So, the amplitude of the prosthesis plate 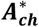, as a function of frequency ***f***_(***i***)_, takes the form the same as the amplitude of the stapes footplate ***A***_***SF***_ in the normal ear

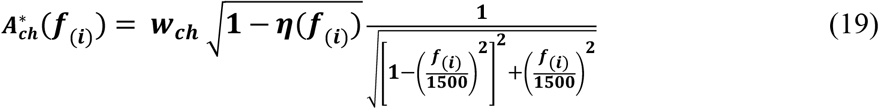

A comparison of the amplitudes 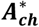 with the tests results ***d***_***ch***_ given in [4] and their relative differences

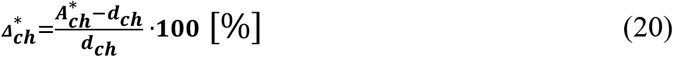

are shown in Table 5. One can see that the absolute value of 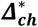 does not exceed **26 %**. Graphs of 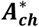 and ***d*_*ch*_** as well as their comparison with 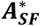 as functions of ***f***_(***i***)_, for **800 *Hz*** ≤ ***f***_(***i***)_ ≤ **8000 *Hz***, based on Eqs (13) and (16) are shown at Fig. 11.

**Table 5.**

| Test (i) | Frequency $f_{(i)}$ [Hz] | Measured amplitude $d_{ch}$ [mm] | Measured amplitude $d_{SF}$ [mm] | Input Ratio IR [dB] | Calculated amplitude $A_{ch}^*$ [mm] | Calculated amplitude $A_{SF}^*$ [mm] | Calculated Input Ratio CIR [dB] |
| --- | --- | --- | --- | --- | --- | --- | --- |
| 1 | 800 | $2.34 \cdot 10^{-5}$ | $2.81 \cdot 10^{-5}$ | -1.59 | $1.88 \cdot 10^{-5}$ | $1.27 \cdot 10^{-5}$ | 3.41 |
| 2 | 1000 | $2.57 \cdot 10^{-5}$ | $1.30 \cdot 10^{-5}$ | 5.92 | $1.90 \cdot 10^{-5}$ | $1.09 \cdot 10^{-5}$ | 4.83 |
| 3 | 1250 | $2.05 \cdot 10^{-5}$ | $7.52 \cdot 10^{-6}$ | 8.71 | $1.71 \cdot 10^{-5}$ | $7.78 \cdot 10^{-6}$ | 6.84 |
| 4 | 1500 | $1.64 \cdot 10^{-5}$ | $6.10 \cdot 10^{-6}$ | 8.60 | $1.42 \cdot 10^{-5}$ | $5.33 \cdot 10^{-6}$ | 8.19 |
| 5 | 2000 | $9.06 \cdot 10^{-6}$ | $4.07 \cdot 10^{-6}$ | 6.95 | $8.80 \cdot 10^{-6}$ | $2.75 \cdot 10^{-6}$ | 10.10 |
| 4 | 2500 | $6.11 \cdot 10^{-6}$ | $2.02 \cdot 10^{-6}$ | 9.61 | $5.33 \cdot 10^{-6}$ | $1.63 \cdot 10^{-6}$ | 10.61 |
| 7 | 3150 | $3.75 \cdot 10^{-6}$ | $6.04 \cdot 10^{-7}$ | 15.85 | $3.15 \cdot 10^{-6}$ | $9.74 \cdot 10^{-7}$ | 10.20 |
| 8 | 4000 | $2.18 \cdot 10^{-6}$ | $6.42 \cdot 10^{-7}$ | 10.62 | $1.78 \cdot 10^{-6}$ | $5.93 \cdot 10^{-7}$ | 9.55 |
| 9 | 5000 | $1.50 \cdot 10^{-6}$ | $7.03 \cdot 10^{-7}$ | 6.58 | $1.16 \cdot 10^{-6}$ | $3.85 \cdot 10^{-7}$ | 9.58 |
| 10 | 6000 | $1.09 \cdot 10^{-6}$ | $5.51 \cdot 10^{-7}$ | 5.93 | $8.37 \cdot 10^{-7}$ | $2.76 \cdot 10^{-7}$ | 9.64 |
| 11 | 8000 | $4.74 \cdot 10^{-7}$ | $1.82 \cdot 10^{-7}$ | 8.31 | $4.94 \cdot 10^{-7}$ | $1.62 \cdot 10^{-7}$ | 9.68 |

**Fig. 11.**
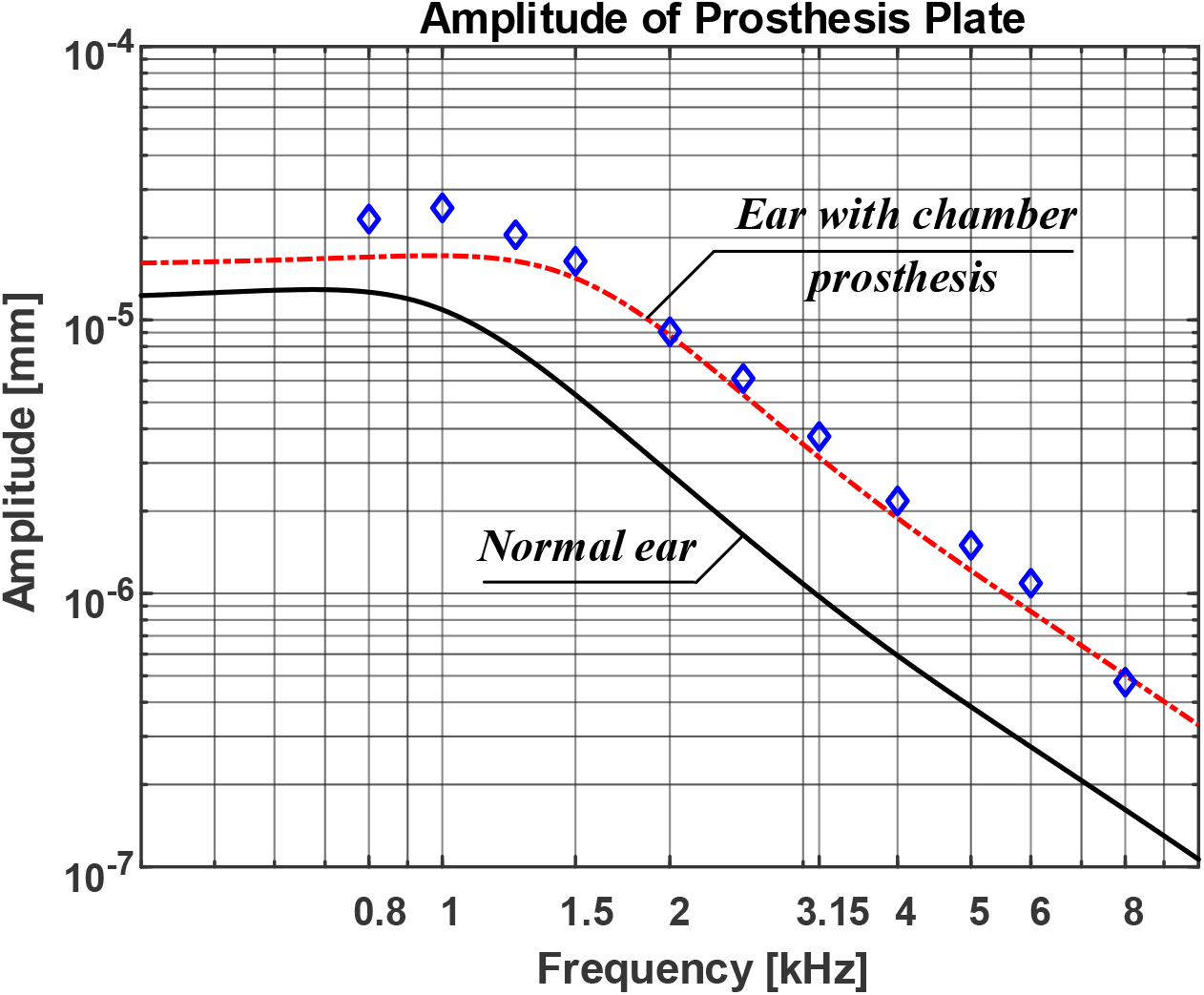
Amplitude of the prosthesis plate: measured ***d***_***ch***_ [4] (diamonds) and calculated 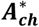 (dashed line); and amplitude of the stapes footplate of the normal ear (solid line).

One can see, that ***k*_*ch*_** has a major impact on the magnitude of 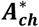. The shift of the resonant frequency to 1500 Hz makes a growth of the amplitudes 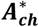 for ***f***_(***i***)_ > **1000 *Hz*** (Fig 9). Anyway, due to lack of tests for frequencies smaller than 800 Hz, there is not a full view of how the ear with the prosthesis behaves at low sound frequencies.

To compare the ear with the chamber prosthesis with the normal ear, the Input Ratio as 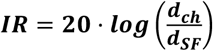, is introduced in the work [4]. In the same way one can introduce a function called Calculated Input Ratio (**c*IR*** [*dB*]) as

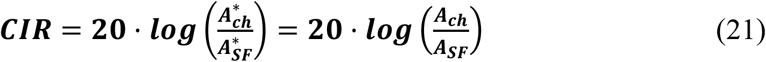

From Eqs (3a, 15) yields that 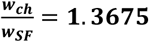. So, based on Eqs (13, 19) one can get the function

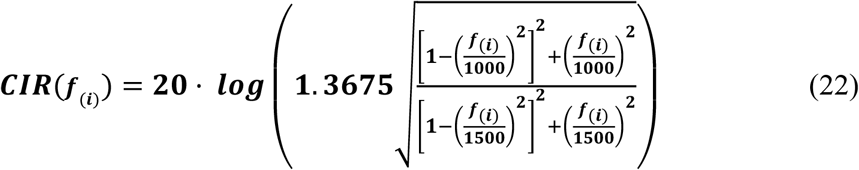

Values of this function for **800 *Hz*** ≤ ***f***_(***i***)_ ≤ **8000 *Hz*** are given in the Table 5, and its graph at Fig. 12. For comparison the values of *IR* given in the work [4] are cited.

**Fig. 12.**
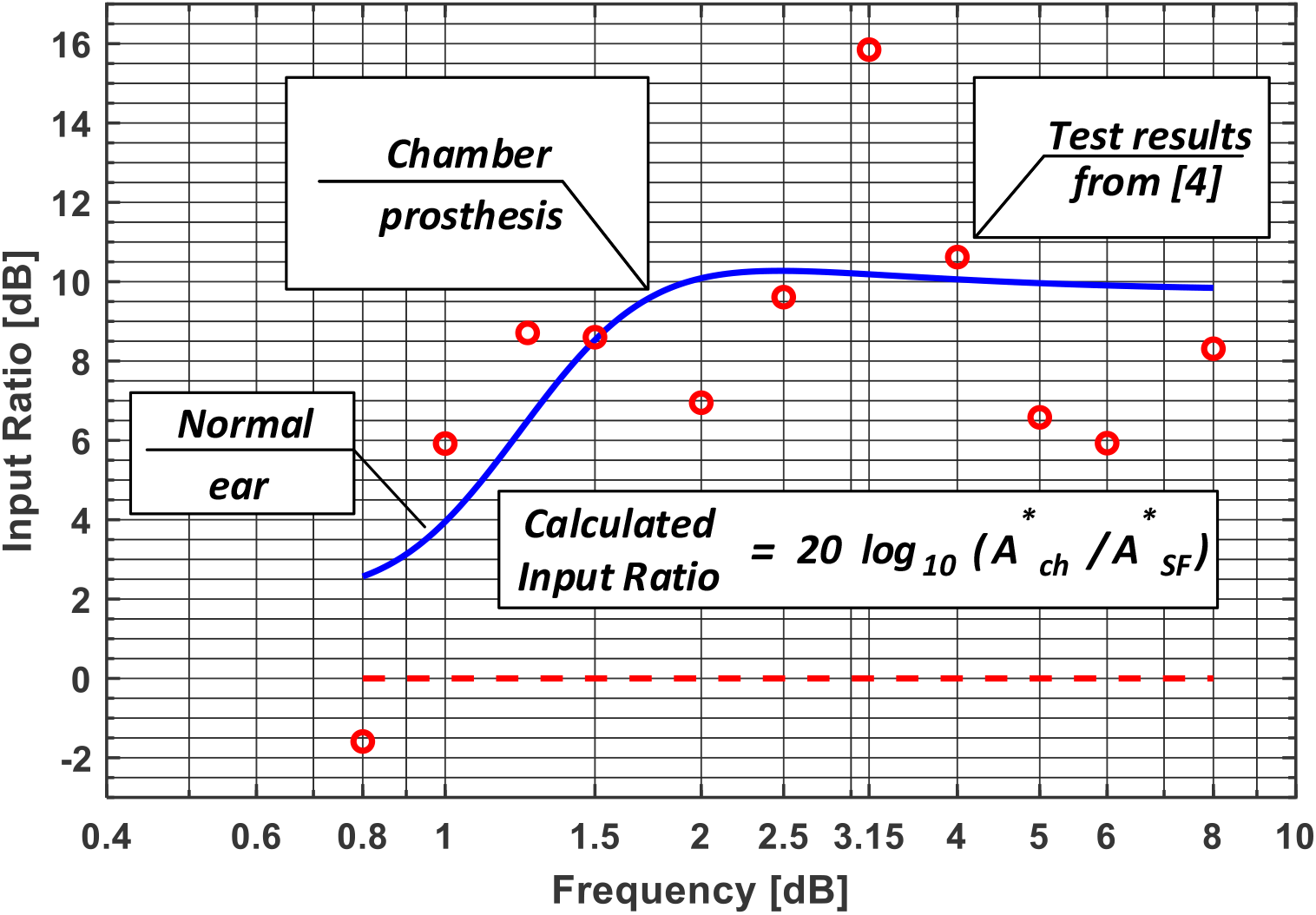
Input Ratio of the chamber prosthesis compared to the normal ear: calculated (solid line), for the normal ear (dashed line), test results (circles).

Note that when ***f***_(***i***)_ ≥ **1000 *Hz***, the Calculated Input Ratio is 4 dB higher for the ear with the chamber prosthesis than for the normal ear. When ***f***_(***i***)_ ≥ **2000 *Hz***, this difference is close to 10 Hz. The test results given in [4] indicate that for ***f***_(***i***)_ ≥ **1000 *Hz*** the Input Ratio may be negative one. It means that the amplitude of the prosthesis plate may be smaller than the amplitude of the stapes footplate in the normal ear. To explain and include to the model this effect, more tests for low sounds are needed.

#### 3.1.3 Flow through the chamber

The vibrating plate of the prosthesis causes not only a sound wave in the fluid filling the chamber, but also a flow of the fluid to the cochlea. The conical shape of the chamber leads to the growth of the velocity of the fluid at its outlet. From the flow continuity equation yields that this growth is given by the ratio of the basis area to the outlet area of the chamber 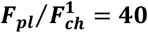. Thus, in the cochlear fluid, a stream appears with a diameter of **0.3 *mm*** and a flow rate **40** times greater than in the rest of the fluid. The stream, hitting the round window membrane, with a diameter of **1.8 *mm***, makes the bulges shown at Fig. 3.

It remains to check, that the 40-fold increase in fluid velocity does not change a laminar nature of the flow. The maximum measured displacement of the prosthesis plate takes place for ***f***_(***i***)_ = **1 *kHz*** and it is ***u*_0_ = 2. 54 10^−5^ *mm*** (see [4]). This corresponds to a velocity of the fluid equal to 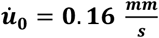. Let ***L*_0_ = 3 *mm*** is the larger diameter of the plate, and 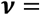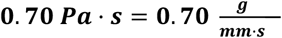 is the kinematic viscosity of the fluid at 36° C. Then the Reynolds number of flow near the plate is ***Re***_**0**_ = ***u***_**0**_***L***_**0**_/***v*** = **0. 6857**. Now, let us check the flow at the outlet of the prosthesis chamber. Now, the fluid velocity is 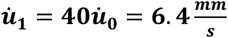 and the corresponding Reynolds number is ***Re***_**1**_ = **40*Re***_**0**_ = **27. 428**. Because the critical value of the Reynolds number is **2300**, the flow through the chamber remains the laminar one.

Note that the narrowing of the chamber causes a growth of flow resistance. The conical shape of the chamber reduces the energy of the fluid flow from the chamber to the cochlea, but it has no influence on the energy of the sound wave. To show it, compare the power of the sound wave in the cochlea ***p***_***c***_ and the power of fluid flow ***p***_***f***_ in the chamber. For the sound intensity *β* = 90 *dB*, the highest measured amplitude of the plate of the prosthesis is ***d***_***ch***_ = **2. 57 · 10**^**−5**^ [***mm***], at the sound frequency ***f***_(***i***)_ = **1000 *Hz*** (see [4]). According to [5], the power of the sound wave ***p***_***c***_ is equal to the power of the prosthesis plate *p*_*ch*_. Taking into account the damping of the round window membrane we have 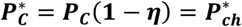 we have (compare Eq. (9))

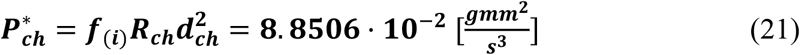

where 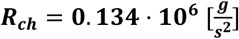 is the stiffness of the plate suspension (see Table 1).

According to the Bernoulli theorem, the power of the fluid flow *p*_*f*_ is the same at each cross section of the chamber. It is sum of the work done in a unit time by the static pressure *p*_*s*_ and the dynamic pressure ***p***_***d***_ of the fluid. If 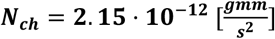 is the maximal amplitude of the force acting on the plate (see Eq. (14)), and ***F*_*ch*_ = 2. 8 [*mm*^2^]** is the plate area, then the static pressure of the plate on the fluid is

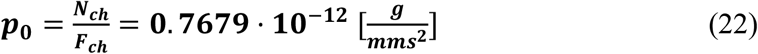

Because the maximal fluid velocity close to the plate is 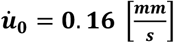, the dynamic pressure close to the plate takes the value

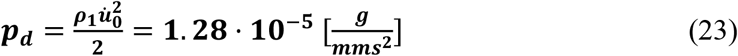

Thereby the power of the fluid in the chamber is

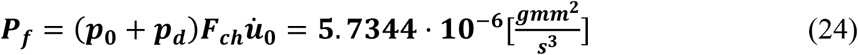

One can see that the ratio of the flow power to the wave power is

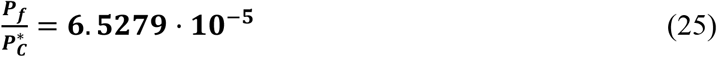

and so, the power of the flow through the chamber is over 10000 smaller than the power of the sound wave. One can state that the flow in the prosthesis chamber has no influence on the sound wave propagation in the cochlea.

## 4. Conclusions

The given in the work [5] model of sound propagation in the human ear enables a simple judge of the use of new implant of the ear. In the new implant the vibrations which create the sound wave in the cochlea take place at the base of conical chamber located in the middle ear niche.

Conclusions which yield from the used model confirm the test results. For sounds higher than 1000Hz, the input amplitude of the sound wave in the ear with the chamber prosthesis is 5-10 dB higher than that in the normal ear. This is due to two facts. The first fact is the shortening of the incus arm which increases the malleus-incus leverage ratio and the force which act on the prosthesis plate. The second is a reduction of the mass of the vibrating plate with the attachment by gluing this element to the incus arm. This makes that the resonant frequency of the plate is higher and the higher sounds are amplified.

It is shown that the flow through the conical chamber has no effect on the amplitude of the input wave in the cochlea. To explain the drop of this amplitude for low sounds needs more tests in this range.

## Footnotes

* According to the project, the mass of the plate is 0.85 mg, the mass of the attachment is 1.72 mg. But after gluing the element to the incus, part of the mass of the attachment is taken by the incus. So, only one third of the attachment mass is taken into account. In the normal ear, the stapes mass is suspended freely on the lenticular process. More on this is given in the section 3.1.2.

** Obtained on the basis of the FE simulation.

